# Live 3D imaging and mapping of shear stresses within tissues using incompressible elastic beads

**DOI:** 10.1101/2021.05.10.443363

**Authors:** Alexandre Souchaud, Arthur Boutillon, Gaëlle Charron, Atef Asnacios, Camille Noûs, Nicolas B. David, François Graner, François Gallet

## Abstract

To investigate the role of mechanical constraints in morphogenesis and development, we develop a pipeline of techniques based on incompressible elastic sensors. These techniques combine the advantages of incompressible liquid droplets, which have been used as precise *in situ* shear stress sensors, and of elastic compressible beads, which are easier to tune and to use. Droplets of a polydimethylsiloxane (PDMS) mix, made fluorescent through specific covalent binding to a rhodamin dye, are produced by a microfluidics device. The elastomer rigidity after polymerization is adjusted to the tissue rigidity. Its mechanical properties are carefully calibrated in situ, for a sensor embedded in a cell aggregate submitted to uniaxial compression. The local shear stress tensor is retrieved from the sensor shape, accurately reconstructed through an active contour method. *In vitro*, within cell aggregates, and *in vivo*, in the prechordal plate of the Zebrafish embryo during gastrulation, our pipeline of techniques demonstrates its efficiency to directly measure the three dimensional shear stress repartition within a tissue, and its time evolution.

## INTRODUCTION

The cohesion and morphogenesis of living tissues require coordinated processes at the cellular scale, based on changes in cell number, size, shape, position and packing (Heisenberg and Bellaïche, 2013; Guirao et al., 2015). These rearrangements are possible because cells can generate and exert mechanical stresses on their surroundings, or conversely feel the stresses and transduce them into biological signals. The complete process is thus regulated under the dual control of genetics and mechanics, which mutually feedback on each other, and drive the growth and shape of tissues (Desprat et al., 2008). Hence, the impact of mechanics on tissue fate and organization is considerable, either for healthy organisms during embryo development (Krieg et al., 2008; Le Goff et al., 2013; Heisenberg and Bellaïche, 2013; Hiramatsu et al., 2013; Hamada, 2015; Herrera-Perez and Karen, 2018), or in pathological conditions (Wells, 2013; Delarue et al., 2014; Angeli and Stylianopoulos, 2016). Quantitative studies about the role of mechanical constraints in morphogenesis and development benefit from a precise and quantitative knowledge of the spatial distribution of mechanical stresses, from the subcellular scale to the tissue scale, and of its temporal evolution.

In the past decades numerous methods have been developed in order to achieve *in situ* stress measurements, using different and complementary techniques; for reviews see Sugimura et al. (2016); Campàs (2016); Roca-Cusachs et al. (2017); Gomez-Gonzalez et al. (2020). To summarize, these techniques can be classified into approximately four categories : (i) External contact manipulations, including micropipettes (Mitchison and Swann, 1954; Hochmuth, 2000; Von Dassow et al., 2010; Guevorkian et al., 2010), microplates (Desprat et al., 2005; Mitrossilis et al., 2009; Tinevez et al., 2009; Mgharbel et al., 2009), AFM indentation (Butt et al., 2005; Elkin et al., 2007; Xiong et al., 2009; Franze, 2011; Lau et al., 2015), traction force microscopy (TFM) (Nier et al., 2016) ; (ii) Manipulations using light, comprising laser ablation (Rauzi et al., 2008; Bonnet et al., 2012; Porazinski et al., 2015) and optical tweezers (Neuman and Nagy, 2008; Bambardekar et al., 2015) - and also by extent magnetic tweezers (Hosu et al., 2003; Tanase et al., 2007; Mazuel et al., 2015) ; (iii) Non-contact optical imaging, in which one can find birefringence (Nienhaus et al., 2009; Schluck and Aegerter, 2010) and stress inference (Chiou et al., 2012; Ishihara et al., 2013; Brodland et al., 2014; Roffay et al., 2021) ; and finally (iv) Embedded local sensors, from FRET at the molecular scale (Grashoff et al., 2010; Borghi et al., 2012) to microsensors at the cell scale (Campàs et al., 2014; Dolega et al., 2017; Mongera et al., 2018; Mohagheghian et al., 2018; Lee et al., 2019; Träber et al., 2019).

The latter technique based on microsensors is quantitative, barely perturbative, and suitable to monitor tissue stresses at the scale of a cell or a group of cells. Two main avenues have already been explored.

The pioneer article (Campàs et al., 2014) has used incompressible liquid droplets to measure the shear stress tensor, which is the most important stress component to understand how anisotropic forces govern tissue morphogenesis. Liquid microdroplets, made of fluorocarbon oil, have been injected in aggregates of mesenchymal cells, in living mandible explants. Coating the oil surface with a biocompatible surfactant enables the droplet insertion in the tissue. The mechanical stresses exerted by the surrounding cells modify the droplet shape, and the deviation from the average stress normal to the droplet surface can be calculated from its local curvature, according to Laplace’s law. Direct and accurate measurements of the three-dimensional (3D) components and orientations of the shear stress tensor require a precise control and calibration of the liquid/tissue surface tension. The same group has refined the technique and successfully applied it to Zebrafish embryos (Mongera et al., 2018).

Other groups have favored elastic beads because they are easier to produce, tune, calibrate, manipulate, insert in tissues, and analyze. The stress exerted on a solid sensor by surrounding cells can be deduced from its deformation, provided that the elastic moduli are determined through an independent calibration. Inspired by Matrigel pressure sensors (Monnier et al., 2016), sensors have been prepared using polyacrylic acid (PAA) hydrogels (Dolega et al., 2017; Lee et al., 2019; Träber et al., 2019), whose Young modulus can be tuned from 60 to 4000 Pa (Lee et al., 2019), or alginate gels (Mohagheghian et al., 2018), and injected in cell aggregates or Zebrafish embryos. Water can flow in and out of an hydrogel, making it compressible. In principle this method yields access to the whole stress, that is, simultaneously the compression stress (including osmotic pressure contributions) and shear stress. The rest state for each sensor without stress can be determined at the end of the experiment by lysing the cells ; once the compression stress is determined, the shear stress can be estimated by substraction (Mohagheghian et al., 2018).

To combine the chemical and mechanical advantages of incompressible liquid droplets, namely fluorescence, functionalization, and accurate shear stress measurements, with the ease to tune and use compressible solid beads, we develop a pipeline of techniques based on incompressible solid beads of diameter comparable to the cell size. The material must exhibit a well-defined elastic behavior, with a Young modulus comparable to the one of the surrounding tissue (order of magnitude 10^3^ Pa), in order to get a measurable deformation under physiological stresses, which expected order of magnitude is e.g. 10^2^ Pa in zebrafish development (Mongera et al., 2018). Coating of the sensors surface is necessary to make them biocompatible and to make their insertion nonperturbative. To observe the sensor’s deformation and get a precise 3D reconstruction of its shape, a stable fluorescent labelling is also needed.

Our choice fell on polydimethylsiloxane (PDMS), which is an elastic elastomer, with a Young modulus adjustable down to a few hundred Pa (Hobbie et al., 2008). Production of small droplets of PDMS polymerisable mixture, having a fixed diameter, can be easily controlled through a microfluidic device. We introduce a novel method to bind the elastomer to a fluorescent dye through covalent bonds, leading to a stable, homogeneous and high intensity fluorescence. Coating the PDMS with cell adhesion proteins is also possible in principle. The sensors can be embedded in the tissue in a non-perturbative way, and their deformation followed over minutes or hours. For 3D image analysis, we implement an active contour method algorithm to determine the shape of the deformed sensors. This method leads to direct and accurate measurements of the 3D components and orientations of the shear stress tensor.

The technique was successfully tested in two different systems, *in vitro* and *in vivo*. First, reconstituted cell aggregates were chosen as a tumor model, for which it is well known that mechanical constraints have a major influence on the organization and fate (Delarue et al., 2014; Northcott et al., 2018). Moreover, it is relatively easy to produce aggregates with embedded sensors, which makes it a privileged system to validate the method. Second, we investigated the distribution of mechanical constraints in the prechordal plate of the Zebrafish embryo during its development. Indeed, based on *in vitro* observation, it has been postulated that anisotropies and heterogeneities of mechanical stresses are present in the prechordal plate, and are of importance to guide its migration (Weber et al., 2012; Behrndt and Heisenberg, 2012). However, due to the lack of appropriate tools, the existence of such anisotropies could not be directly tested so far. The implantation of our mechanical sensors in this system could definitively help to decide between different models actually disputed.

In both cases, *in vitro* and *in vivo*, we report here results concerning the spatial repartition and temporal evolution of the shear stresses, providing clear demonstration of the usability and potential of these new sensors.

## METHODS

The above requirements can be summarized as follows : the sensors must have a size comparable to the cell size, and be easily fabricated and manipulated in large quantities, with monodisperse radius ; they must be biocompatible and conveniently embedded in living tissues ; their rigidity has to be homogeneous and close to those of the tissues ; they can be fluorescently labelled in a homogeneous and stable way ; their 3D shape can be precisely reconstructed with minimal effort ; the effective shear modulus of the sensors can be reliably calibrated *in situ* ; the shear stress tensor can be decoupled from compression stress and directly derived from the 3D sensor’s deformations by using linear elasticity ; shear stresses of order of 10^2^ Pa can be measured with 10 Pa sensitivity. To satisfy simultaneously all these constraints, we introduce the following pipeline of techniques.

### Microsensors fabrication

The microsensors were made out of a silicon elastomer similar to usual PDMS. The polymerizable mix preparation is first dispersed into liquid droplets of about 30 µm in size, thanks to a microfluidic circuit, and afterwards polymerized at 80 ±1 °C. Briefly, the main component of the elastomer is vinyl terminated polydimethylsiloxane, hereafter coined DMS. It is mixed with a polymeric hydrosilane (methylhydrosiloxane-dimethylsiloxane copolymer) that acts as a cross-linker via hydrosilylation of the vinyl ends of DMS (Fig. S1a). The ratio of cross-linker to DMS must be carefully controlled (*mcross* = 1.60 ×10^−2^ *mDMS*) to achieve the desired shear modulus after polymerisation. The hydrosilylation reaction is initiated using Karstedt’s catalyst (*m*_*catal*_ = 4.286 × 10^−3^ *m*_*DMS*_). A divinylic inhibitor (diallyl maleate) is also added (*m*_*inhib*_ = 0.8571 × 10−3 *m*_*DMS*_) to slow down the reaction kinetics. The as-prepared mixture needs to be stored at 4°C and should be used within a few days least the cross-linking reaction significantly moves forward.

Dispersing the polymerizable mix into spherical droplets of homogeneous size is obtained through a custom-made microfluidic circuit, by a classical flow focusing method (Haejune, 2007). The dispersed phase (polymer mix) meets the carrier phase (water) at a 4-channels crossing, and the resulting droplets suspension is collected at the output. The rectangular channel section is about 50 × 20 µm^2^. The injecting pressures of both phases is finely tuned and regulated by a Fluigent controller, in order to get a steady-state dripping instability and a constant droplet diameter. This diameter may be tuned between 20 and 40 µm. It is stable for a given droplet batch : the diameter distribution of droplets is quite monodisperse within a same production (Fig. 1).

**Fig. 1.**
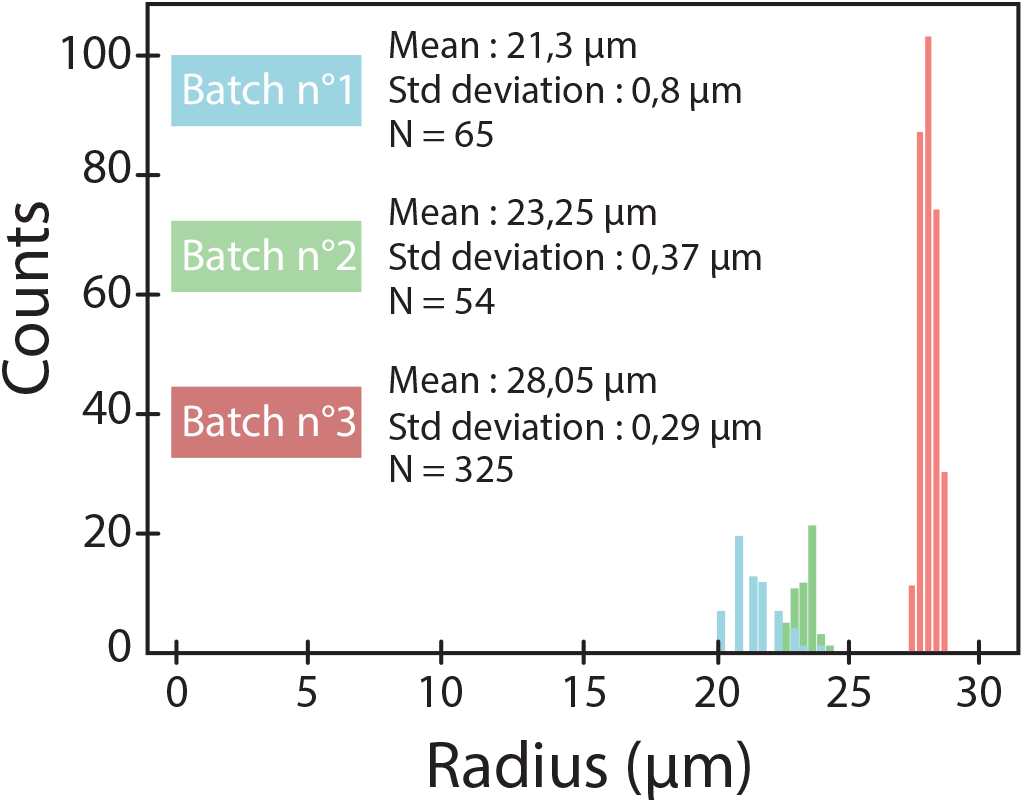
Distribution of microsphere’s radii for three different batches obtained with a microfluidic device

The droplets polymerization into spherical elastic beads is achieved by baking the suspension at 80°C during 3 h. The final gel is very soft. It sticks easily and irreversibly to any wall, including inert surfaces like teflon or silanized glass. Thus one must avoid the contact of the droplets with any solid surface during and after polymerization. To achieve this, the beaker containing the suspension is placed during baking on a turntable, rotating at about one turn per second. Since the gel is less dense than water, buoyancy makes the polymerized beads spontaneously concentrate at the center of the meniscus of the water free surface. Then, the concentrated suspension containing about 10^4^ beads per mL can be collected with a micropipette, aliquoted in Eppendorf tubes and immediately stored in a freezer at − 20°C. The beads remain stable for weeks at this temperature, and are thawed before immediate use.

### Fluorescent labelling

In order to observe the sensors embedded in the tissue, a dye must be added to the elastomer. The objective is to get an homogeneous fluorescent signal, with high enough intensity, to be able to visualize the sensors with usual fluorescent microscopy techniques (confocal, spinning-disk and 2-photon microscopy) and to precisely reconstruct their 3D shape. Several hydrophobic dyes did not lead to satisfying labelling : either the dye could not be homogeneously dispersed in the polymer mix (fluorescein diacetate), or it was partially released in the water solution surrounding the beads, so that the fluorescent signal rapidly decreased with time (Nile Red and Cryptolyte™). We also attempted to label the elastomer with quantum dots (QDs) dispersed in the mixture. However, despite some specific coating to make them hydrophobic, QDs remained partially aggregated and the dispersion was not complete.

For efficient fluorescent labelling of the elastomer, two challenges needed to be overcome: the dye had to easily disperse into the polymerisable mix and, once dispersed and after curing, should not escape the meshwork of the gel and leak into the surroundings. Our strategy entailed attachment of the organic dye to the cross-linker via a parallel hydrosilylation reaction (Fig. S1b). The dye therefore needed to bear a vinyl terminal group. Isothiocyanate-bearing fluorophores can be conveniently modified through quantitative C-N bond formation. We selected Rhodamine B isothiocyanate because of its emission in the red and condensed it with allylamine to give a vinyl-terminated rhodamine B analogue (Fig. S1c). The compound was then added to DMS in a molar ratio of one fluorophore for 1,000 DMS strands (see Supplementary Files for details).

Figure 2 shows some examples of bright field and fluorescent images of single sensors (a) suspended in water ; (b) embedded in a CT26 reconstituted aggregate ; (c) implanted in a zebrafish embryo. Image (b2) is a 3D reconstruction obtained with ImageJ software, where the deformation from spherical shape is clearly visible. The fluorescent signal is homogeneous in the bead volume and the contrast is high enough to detect the sensor border (see section **Active contour method**). Moreover, we checked that the fluorescence in the elastomer remains stable over several days. Thus this labelling technique fulfills all the requirements for further quantitative image analysis.

**Fig. 2.**
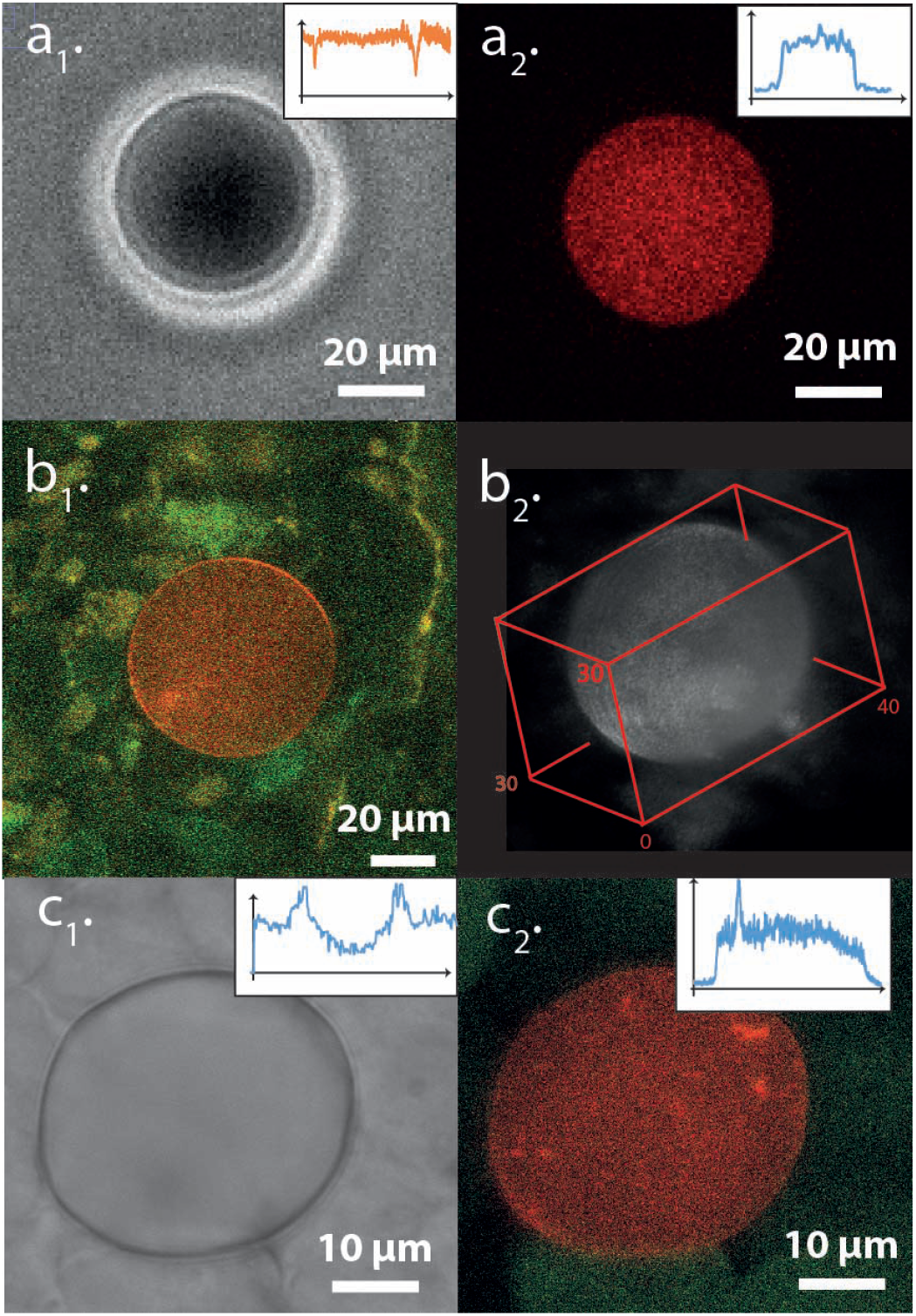
Images of three sensors in different situations : (a) suspended in water ; (b) embedded in a CT26 reconstituted aggregate (green) ; (c) implanted in a zebrafish embryo (green). (a1) and (c1) are bright field images. (a2) is obtained with a spinning-disk, (b1) with a 2-photon and (c2) with a confocal microscope ; (b2) is a 3D reconstruction obtained with ImageJ software. The insets represent the intensity profile through the sensor’s diameter

### Strain-stress relationship

When a bead is embedded in a tissue, it experiences mechanical forces exerted by its environment, which induce a deformation from its initial spherical shape, and makes it a sensor of local stresses. In the following we assume that the sensor is small enough, as compared to the length scale characterizing stress spatial variations, so that the stress tensor can be considered as homogeneous over the sensor’s volume. Thus, stress variations at a scale smaller than the sensor cannot be detected. We also assume that the elastomer shows an ideally elastic behavior, and that its deformation remains small enough (≲ 5%), so that linear elasticity applies. The local strain tensor 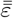 and stress tensor 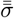 are related through (Landau et al., 1986):

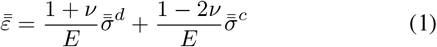

Here 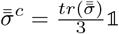 represents the isotropic part of 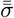 (traction stress tensor, equivalent to a pressure) and 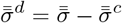 is the deviator, also known as shear stress tensor ; *E* is the Young modulus, and *ν* the Poisson’s ratio.

The sensors are made out of a PDMS elastomer, which can be considered as incompressible in the range of physiological stresses (its compression modulus 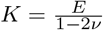 is of the order of 10^6^ Pa). This justifies the approximation 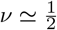, and thus Eq. 1 simplifies into : 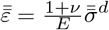. Introducing the shear modulus 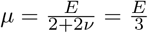, one gets:

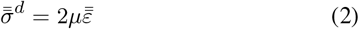

which we will use in the following.

Under external stresses, and assuming small deformations, the sensor’s shape changes from a sphere of radius *a* to an ellipsoid of half-axes *a*_*X*_, *a*_*Y*_ and *a*_*Z*_. In the system of principal coordinates (*X, Y, Z*) of this ellipsoid, both the strain and stress tensors are diagonal, so that one can write :

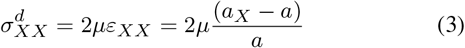

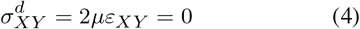

and similar relations for other stress components. Hence, provided that the shear modulus of the PDMS elastomer has been independently calibrated, the local shear stress tensor is fully determined by pointing the ellipsoid orientation and measuring the length of its half-axes.

### Cell culture and preparation of aggregates

CT26 cells, stably transfected with Lifeact-GFP, were cultivated in T75 flasks at 37°C in 5% CO_2_, in DMEM culture medium completed with 10% (v/v) Foetal Bovine Serum and 1% antibiotics (penicillin-streptomycin), and passed every 3 days. For the preparation of aggregates, confluent cells were detached by using 5 mL of buffer solution containing 0.05% trypsin. Incubation was limited to about 1 min, in order to form cell leaflets and avoid complete cell dispersion. In parallel, elastomer microsensors were functionalized by adding 1 mL of fibronectin solution in PBS (50 µg*/*mL) to 1 mL of freshly thawed bead suspension. The final suspension was left for incubation during 1 h at 37°C, and then directly added to ∼ 10 mL of detached cells suspension without further rinsing. Aggregates containing inserted beads were prepared in Petri dishes on which the cell and bead suspension was deposited, placed on an orbital agitator (∼50 rotations*/*min), and left to grow for at least 24 h in an incubator at 37°C. To get spherical aggregates, it is suitable to let them grow for at least 48 h. The diameter of the obtained aggregates lies between 100 and 500 µm. Within few exceptions, they contain at most one bead per aggregate.

### Zebrafish preparation

Embryos were obtained by natural spawning of *Tg(−1*.*8 gsc:GFP)ml1* adult fishes (Doitsidou et al., 2002). All animal studies were approved by the Ethical Committee N°59 and the Ministère de l’Education Nationale, de l’Enseignement Supérieur et de la Recherche under the file number APAFIS#15859-2018051710341011v3.

Embryos were grown at 28.5°C until reaching shield stage (6 hours post fertilization). Embryos were then processed as explained in Boutillon et al. (2018). Using a large glass needle (35 µm opening) mounted on a pneumatic microinjector (Narishige IM-11-2) under a fluorescence-stereo microscope, a sensor was inserted in the shield of an embryo, which expresses GFP in the *Tg(−1*.*8 gsc:GFP)ml1* line. Transplanted embryos were then incubated at 28.5*°C until* reaching the desired stage between 60% and 85% of epiboly (6,5 to 8 hours post fertilization). Embryos were then selected for the presence of the sensor in the prechordal plate. Selected embryos were mounted in 0,2% agarose in embryo medium on the glass coverslip of a MatTek petri dish (Boutillon et al., 2018) and placed on an inverted TCS SP8 confocal microscope (Leica SP8) equipped with an environmental chamber (Life Imaging Services) at 28°C and a HC PL APO 40x/1.10 W CS2 objective (Leica). Imaging parameters were set to acquire the whole sensor (*z*-stack) in less than 15 seconds, to minimize displacement due to the migrating neighboring cells.

### Image recording and analysis

#### Microscopy

To simultaneously image the tissue and the sensors embedded inside it, several techniques have been used:

i. Frequently, sensors in suspension in water were imaged with a spinning-disk microscope (Andor Revolution CSU X1, mounted on an Olympus IX 81 inverted microscope equiped with a 40x water immersion objective), in order to check their sphericity, and the good quality of their fluorescence (intensity and homogeneity) (Fig. 2a).
ii. 2-photon microscopy was used for the visualization of reconstituted cell aggregates. Experiments were carried out at the multiphoton facility of the ImagoSeine imaging platform (Institut Jacques Monod, Université de Paris). The aggregates were deposited in a Petri dish, in a chamber regulated at 37°C, and observed during a few hours under a 20x water immersion objective at the early stage of their adhesion to the bottom plate. For the rhodamin dye, the excitation laser was tuned at *λ* = 840 nm and the emitted light was collected through a dichroïc mirror at *λ* ≥ 585 nm. Lifeact-GFP of CT26 cells was excited at *λ* = 900 nm and the fluorescence was collected at *λ* ≤ 585 nm. Image stacks were recorded along the optical axis every 0.5 µm, with a lateral resolution down to 0.1 µm*/*pixel (Fig. 2b).
iii. A confocal microscope (Leica SP8) was used to image the prechordal plate of the Zebrafish embryos. The sample was maintained at *T = 28°C*. Image stacks (x40 water immersion objective) were recorded at regular time intervals (30 s to 1 min) at different stages of the epiboly, comprised between 60 and 85%. Since the prechordal plate is migrating at a velocity up to 2 µm*/* min, the acquisition time for a whole stack must be smaller than 15 s to avoid drift in the images. Hence images were recorded every 2 µm. The excitation laser was tuned at *λ* = 498 and 550 nm and the emitted light was collected between 507 − 537 nm and 569 − 673 nm (Fig. 2c).

#### Active contour method

A careful 3D reconstruction of the sensor’s shape was required to retrieve the orientation and half-axes of the deformed beads with a good accuracy. Indeed, the usual built-in applications for 3D reconstructions, such as ImageJ plugins, do not lead to a reliable and accurate enough profile: the result depends on specific choices of parameters for the filters and for the intensity thresholds, which involve the subjective appreciation of the operator. Hence, we implemented an active contour method, as follows (Kass et al., 1988; Caselles et al., 1993; Marquez-Neila et al., 2014; Bendaoud, 2017) : the common principle of the different existing algorithms consists in considering a swelling (or shrinking) surface *ν*(*s, n*) at the *n*^*th*^ iterative stage, parametrized by its local coordinates *s* = (*s*_1_, *s*_2_). A function *E*(*ν*) is associated to this surface and, like an energy, is built to reach a minimum when the surface *ν*(*s, n*) coïncides with the contour of the object. This pseudo-energy is the sum of three contributions:

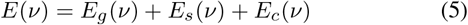

The first term *E*_*g*_ (*ν*) is a gradient detection term :

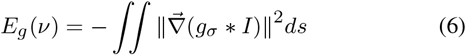

It represents the norm of the intensity gradient, convoluted by a gaussian filter *g*_*σ*_, and integrated over the surface *ν*. The minus sign ensures that *E*_*g*_ (*ν*) has a minimum when the intensity gradient on the surface *ν* is maximal.

The second term (surface energy) *E*_*s*_(*ν*), is analogous to a Helfrich energy (Helfrich, 1973) :

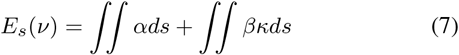

where *α* is a surface tension, *κ* the local curvature of the surface *ν*, and *β* a curvature stiffness. This term limits the roughness of the final contour.

The last term (balloon energy) *E*_*b*_(*ν*) is proportional to the volume *V* limited by *ν*, and forces the surface to swell or to shrink when iterating the process, according to the sign of the parameter *δ* :

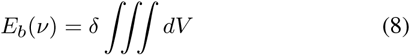

The details of the used python code can be found in (Souchaud, 2020) and on the Github platform (see **Data availability**). Starting from a seed located inside the contour to be detected, and taking *δ <* 0, the volume delimited by *ν* enlarges at each step *n* of the algorithm, until *E*(*ν*) reaches a minimum, which defines the contour of the object. The principle of the method is illustrated in Figure 3a, and a example of contour determination for a micro-sensor is shown in Figure 3b. A movie (M1) showing the 3D reconstruction of a bead inserted in the prechordal plate of a Zebrafish embryo is available in supplementary files. Consequently, the final contour position only depends on the choice of *δ* and of two parameters *α*_1_ and *β*_1_ derived from *α* and *β*. In the algorithm, *δ* is an integer and must be equal to − 1 to ensure convergence. We have checked that tuning *α*_1_ and *β*_1_ in a large range (variations up to 100%) changes the contour position by less than 0.2 µm. Thus the final accuracy Δ on the contour determination is not limited by the algorithm, but by the quality of the image. It is about 0.5 µm, over a sensor radius of about 15 µm.

**Fig. 3.**
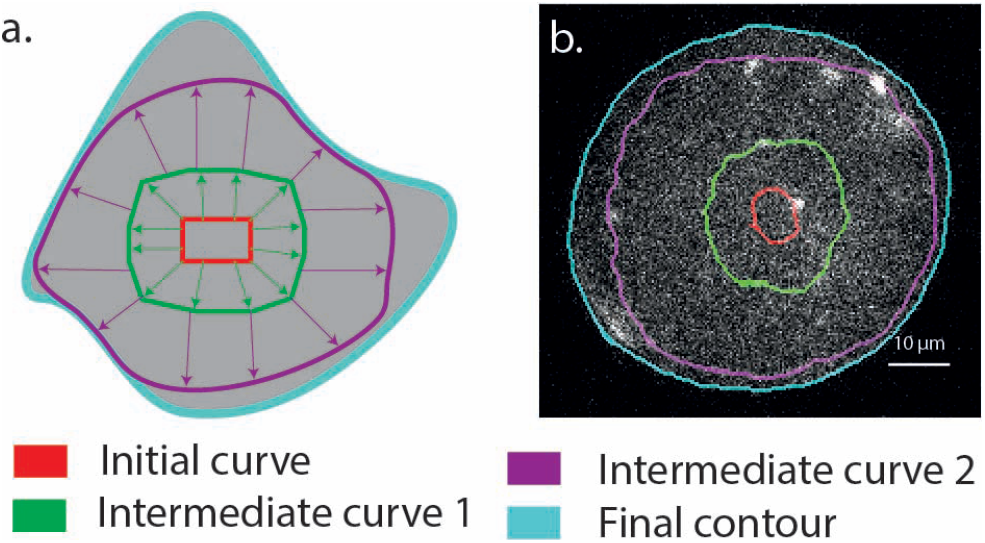
Determination of the sensor’s contour ; (a) Principle of the active contour method : the initial seed (red) progressively swells (green, purple), until it reaches the contour of the object (cyan), which minimizes its pseudo-energy (see text). (b) Example of contour determination for a sensor inserted in an aggregate (same color code as (a)).

Once the 3D contour of the sensor has been determined from the images, a renormalization factor *r* = 0.935 must be applied to the sensor’s shape along the optical axis (*z*) direction. This factor takes into account a geometrical correction due to light refraction through the tissue/PDMS interface, which acts like a spherical diopter between two media of different optical indices (1.35 *< n*_1_ *<* 1.40 for the living tissue ; *n*_2_ ≃ 1.44 for PDMS). The factor *r* was calibrated *in situ*, by comparing the shape of hard non-deformable spherical sensors (*µ* ∼ 2 × 10^4^ Pa) to their reconstructed image. Calibrations were performed both in cell aggregates and in zebrafish embryos, leading to the same value *r* = 0.935 ± 0.02, which was retained in the following.

With this active contour method, we estimate that we can detect a sensor deformation if the difference between two half axes is at least equal to threshold Δ = 0.5 µm, which represents the accuracy of our measurements.

## RESULTS

### Sensors mechanical calibration

As shown in section **Strain-stress relationship**, determining *in situ* the local stress tensor requires to calibrate the sensors shear modulus *µ*. Its value was determined by two methods, first at the macroscopic scale with a commercial rheometer, and secondly *in situ* at the sensor’s scale with a custom-made setup allowing uniaxial compression of aggregates.

#### Macroscopic rheometry

A rheometer (ARES G2) was used to follow the evolution of the elastic moduli of the PDMS preparation during its gelification. The polymerizable mixture was introduced in either plate-plate or cone-plate geometry, and maintained at a constant temperature *T* = 60°C or *T* = 80°C, whilst the storage and loss moduli *G*′ and *G*″ were measured every 15 min, in the range 0.1 Hz *< f <* 10 Hz. After a transient increase during about one hour, *G*′ and *G*″ gradually tend towards a plateau, which final value is reached after ∼ 12 h at *T* = 60°C, or ∼ 3 h at *T* = 80°C.

At any given stage of gelification, *G*′ was found independent of the excitation frequency *f*, and *G*″ increased approximately linearly with *f*, which corresponds to a Kelvin-Voigt behavior. Moreover, at the end of the gelification plateau, the ratio *G′/G″* was found of the order of 10 at *f* = 10 Hz for a standard gel composition. Thus, when the PDMS gel is submitted to a static (or very slowly varying) stress, it may be considered as a purely elastic solid, and it is legitimate to confound its static shear modulus *µ* with its storage modulus *G*′ extrapolated at *f* = 0 Hz.

The *G*′ value, measured at the end of the polymerization plateau, was retained in the following as the value of the shear modulus for bulk PDMS *µ*_*b*_. This value strongly depends on the mixture composition. It is close to 500 Pa for *m*_*cross*_ = 0.0160 *m*_*DMS*_, but reaches 1000 Pa for *m*_*cross*_ = 0.0170 *m*_*DMS*_. We noticed that the final value of *µ*_*b*_ depends also, but in a lesser extent, on the crosslinker and inhibitor concentrations, and on the gelification temperature.

The shear modulus *µ*_*d*_, measured after the polymerization of a mix dispersed in water, appeared to be different from the bulk shear modulus *µ*_*b*_. A small amount of polymerizing mixture was added to water and vigorously shaken for a few seconds to make a coarse emulsion. The suspension was left to buoy up at room temperature, during a timelapse *τ*, after which the suspension was centrifugated until complete droplets coalescence, and the supernatent was sampled and placed in the rheometer. The actual *µ*_*d*_ was always found smaller that the bulk *µ*_*b*_ measured for the same mix before emulsification. The ratio *µ*_*d*_*/µ*_*b*_ reached a stable plateau value ≃ 0.43 when *τ* ≳ 24 h. We interpreted this observation by assuming that a small amount of a mix component, likely the crosslinker, may diffuse out of the DMS emulsion droplets and dissolve in the surrounding water. As seen above, a small variation of the crosslinker concentration is enough to induce a significant change in the final mechanical properties of the gel. Since the microsensors are made from small droplets suspended in water before polymerization, this effect has to be considered for a proper calibration of their mechanical properties. In practice, we decided to measure *µ*_*b*_ with the rheometer for every batch of bulk mix polymer used to make microsensors, and then to apply a constant corrective factor in order to get an estimate of the final shear modulus *µ*_*d*_ of the spherical elastic sensors : *µ*_*d*_ = 0.43 *µ*_*b*_.

#### *In situ* calibration in aggregates

At the microsensor’s length scale, one expects that capillary effects, due to the non-zero surface tension *γ*_*c*_ ∼10 mN*/*m between the tissue and the PDMS sensor of radius *a* ∼15 µm, might affect its global mechanical response (Style et al., 2017; Bico et al., 2018). Indeed, the contribution of the Laplace term 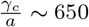 Pa is comparable to the macroscopic shear modulus *µ*_*d*_ of the PDMS dispersion. This means that the relationship between the applied external stress and the deformation of the sensor involves both the shear stress modulus *µ*_*d*_ and the surface tension *γ*_*c*_. For small deformations, it has been shown that one can take into account this elasto-capillary contribution by introducing an effective elastic constant *µ*_*e*_ (Carbonaro et al., 2020):

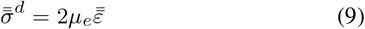

with

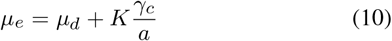

*K* is a dimensionless constant of order unity. We thus performed an *in situ* calibration on a sensor embedded in a tissue, in order to directly measure *µ*_*e*_.

For this calibration, we used a custom-made uniaxial rheometer, allowing to apply either a controlled force, or a controlled deformation to a cell aggregate (Desprat et al., 2005; 2006; Mitrossilis et al., 2010). To summarize, the cell aggregate may be squeezed between two glass plates, a rigid one and a flexible one acting like a cantilever. The plates are actuated by two piezoelectric stages. A feedback loop maintains the extremity of the flexible plate, on the aggregate side, at a fixed position, while its other extremity is free to relax with time. This allows to record the evolution of the force *F* (*t*) exerted on the aggregate, at constant aggregate’s deformation. *F* (*t*) is calculated from the flexible plate’s deflection, knowing its rigidity *k* = 81.2 nN*/*µm.

Practically, we selected a CT26 aggregate of diameter comprised between 100 and 200 µm, containing a sensor localized close to the aggregate center, and we seized it between the two plates of the rheometer (Figure 4a). We then applied a step motion to the rigid plate to squeeze the aggregate, while the flexible plate extremity close to the aggregate is regulated at constant position. From this initial instant we recorded the relaxation of the flexible plate’s deflection during 15 to 30 min, and thus the time evolution *F* (*t*), while the aggregate deformation remained constant. Simultaneously, we imaged the shape of the sensor in its median plane (Figure 4b). Two or three successive squeezings and relaxations were operated on the same aggregate. From these relaxations we inferred at any time *t* the force *F* (*t*) exerted on the aggregate, and the deformations *ε*_*zz*_ (*t*) and *ε*_*rr*_(*t*) of the sensor, respectively in the compression direction and perpendicular to it.

**Fig. 4.**
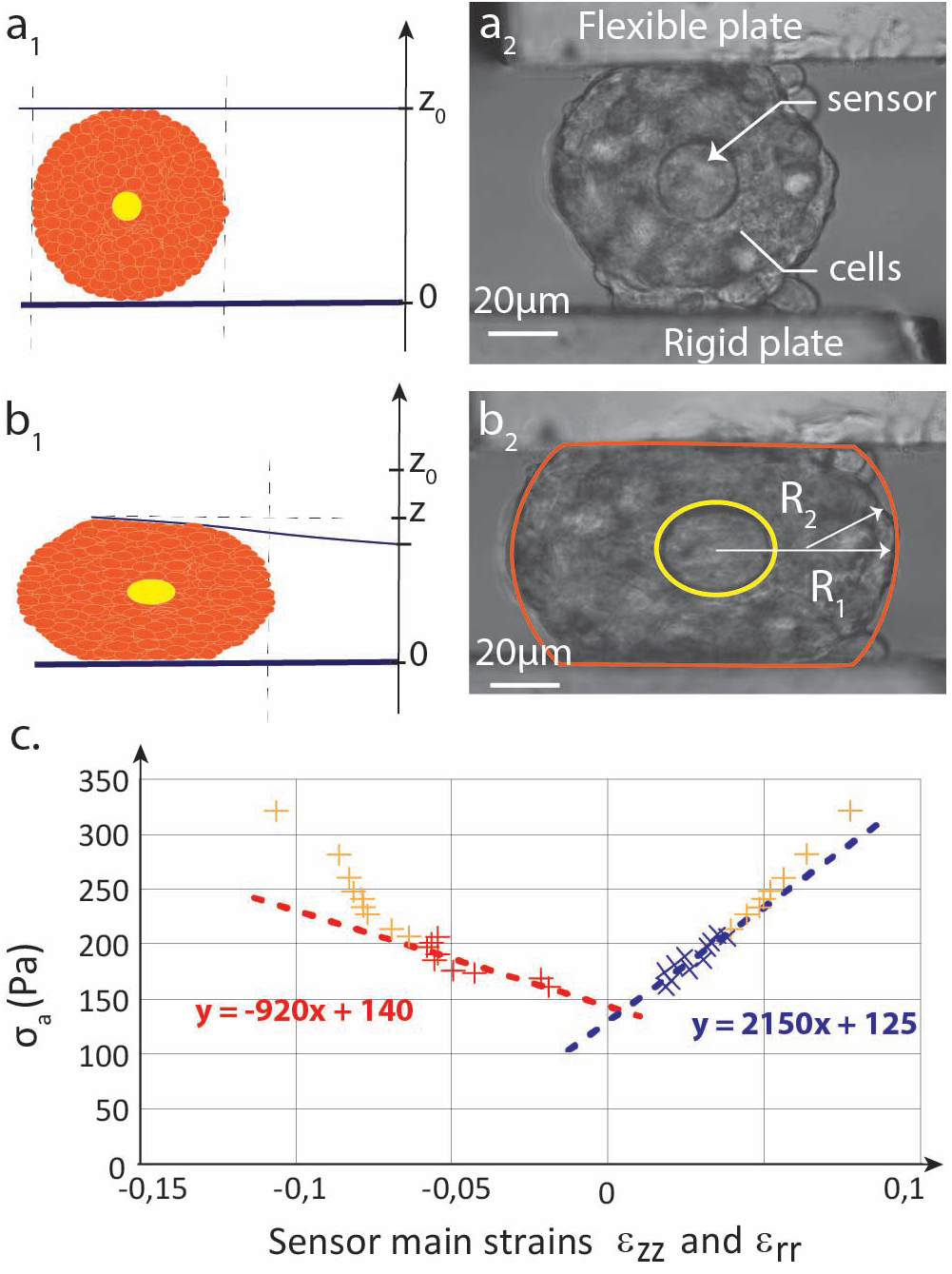
In situ calibration of a sensor shear modulus. Principle (a1, b1) : a CT26 aggregate, initially spherical and containing a sensor at its center, is squeezed between two glass plates. The images (a2, b2) are analyzed to extract the sensor main strains *ε*_*zz*_ and *ε*_*rr*_. (c) Variations of the average stress *σ*_*a*_(*t*) *vs ε*_*zz*_ (*t*) (red crosses) and *ε*_*rr*_ (*t*) (blue crosses) during the relaxation of a squeezed aggregate. Yellow points are recorded during the first ≈ 30 s of the relaxation. The shear modulus *µ*_*e*_ of the sensor can be extracted from the slopes of the straight lines (Eq.12 and **??**).

As detailed in the appendix, we developed a model to establish the relationship between the sensor’s deformation 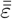, and the average stress in the equatorial plane of the aggregate, defined as :

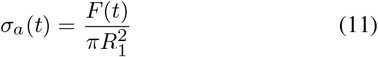

This model predicts :

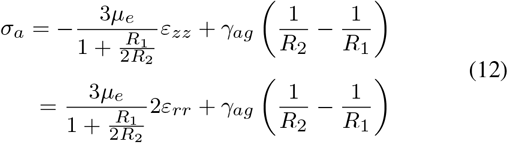

Here *R*_1_ represents the equatorial radius of the aggregate, *R*_2_ its curvature radius in the observation plane (Fig. 4*b*_1_), and *γ*_*ag*_ is the surface tension between the aggregate and the culture medium (not to be confused with the sensor/aggregate surface tension *γ*_*c*_). Eq. (12) is valid under the following approximations : (i) the aggregate is supposed spherical at rest and the *z* axis is a cylindrical symmetry axis at any time ; (ii) the sensor does not perturbate the stress distribution in the aggregate ; (iii) the sensor is approximately at the center of the aggregate ; (iv) the component *σ*_*zz*_ (*t*) is assumed to be homogeneous in any plane perpendicular to the main compression axis.

Figure 4c shows an example of such a stress-strain relationship, measured during the relaxation of a squeezed aggregate. We experimentally check that *σ*_*a*_(*t*) linearly varies with *ε*_*zz*_ (*t*) and with *ε*_*rr*_(*t*) during the relaxation, except for the yellow points which are recorded during the first ≈ 30 sec of the relaxation. Indeed, immediately after the step compression, the stress components quickly vary, due to different relaxation mechanisms in the aggregate. In this initial non-linear regime, it is likely that hypothesis (iii), namely in-plane spatial homogeneity of *σ*_*zz*_ (*t*), is not valid. On the other hand, at longer time, the relaxation slows down and one expects that the homogeneity assumption becomes verified. In this regime, the experimental data meet the model prediction given by Eq. (12). Hence, from the linear fits shown in figure 4c, one can extract the values of the effective shear modulus *µ*_*e*_ = 790 ± 160 Pa and of the aggregate surface tension *γ*_*ag*_ = 9 ± 2 mN*/*m for this particular experiment. Note that we experimentally measure *ε*_*zz*_ ≃ − 2*ε*_*rr*_ at any time, which is consistent with the sensor’s incompressibility and with the cylindrical symmetry assumptions.

Two gels of slightly different compositions have been tested. Within our experimental accuracy, no significant difference can be detected between the value of 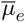, averaged over *N* assays in different aggregates, and the value of *µ*_*d*_ measured at the macroscopic scale for a coarse emulsion made out of the same gel (see Table 1). These results do not allow us to isolate the contribution of capillary effects in the effective shear modulus *µ*_*e*_, according to Eq. 10. Either this contribution is smaller than expected, or the determination of *µ*_*d*_ and *µ*_*e*_ is not accurate enough to measure the difference between them. In the following, we will take *µ*_*d*_ as the reference value for the effective elastic shear modulus of the sensors.

**Table 1.**
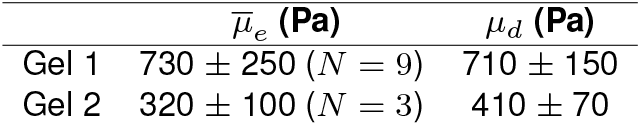
Comparison of the average value 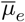, measured *in situ*, and *µ*_*d*_, measured for a coarse emulsion of the same gel. Experiments have been performed on two gels having slightly different compositions.

| | $\bar{\mu}_e$ (Pa) | $\mu_d$ (Pa) |
| --- | --- | --- |
| Gel 1 | $730 \pm 250$ ($N = 9$) | $710 \pm 150$ |
| Gel 2 | $320 \pm 100$ ($N = 3$) | $410 \pm 70$ |

From the uniaxial compression of aggregates we can also infer the surface tension *γ*_*ag*_ between the aggregate and the culture medium. The values range from 3 to 12 mN*/*m for different aggregates. Although the dispersion is important, the order of magnitude corresponds to the expected one.

### Stress distribution in cell aggregates

13 aggregates, containing deformable sensors located at different positions, were imaged with a 2-photon microscope. We analysed the shape of 17 sensors. To compare the results, we define a dimensionless position *r* = *r*_*c*_*/R*_*a*_ as the ratio of the distance *r*_*c*_ from the aggregate center to the sensor center, over the distance *R*_*a*_ from the aggregate center to the aggregate edge in the direction of the sensor. This definition takes into account the fact that the aggregate might be not spherical but slightly ellipsoïdal.

The results are gathered in Fig. 5a. Each sensor is set at its reduced position *r*, and is represented by an ellipse showing its deformation projected in the (*x, y*) plane of the image. Since the actual deformations are small (*<* 10%), they were artificially multiplied by a factor 4 on the scheme to be visible. The main components of the associated shear stress are represented as red bars of length proportional to the stress amplitude. In most cases, one of the main axes remains close to the *Oz* optical axis, which justifies the projection in the (*x, y*) plane.

**Fig. 5.**
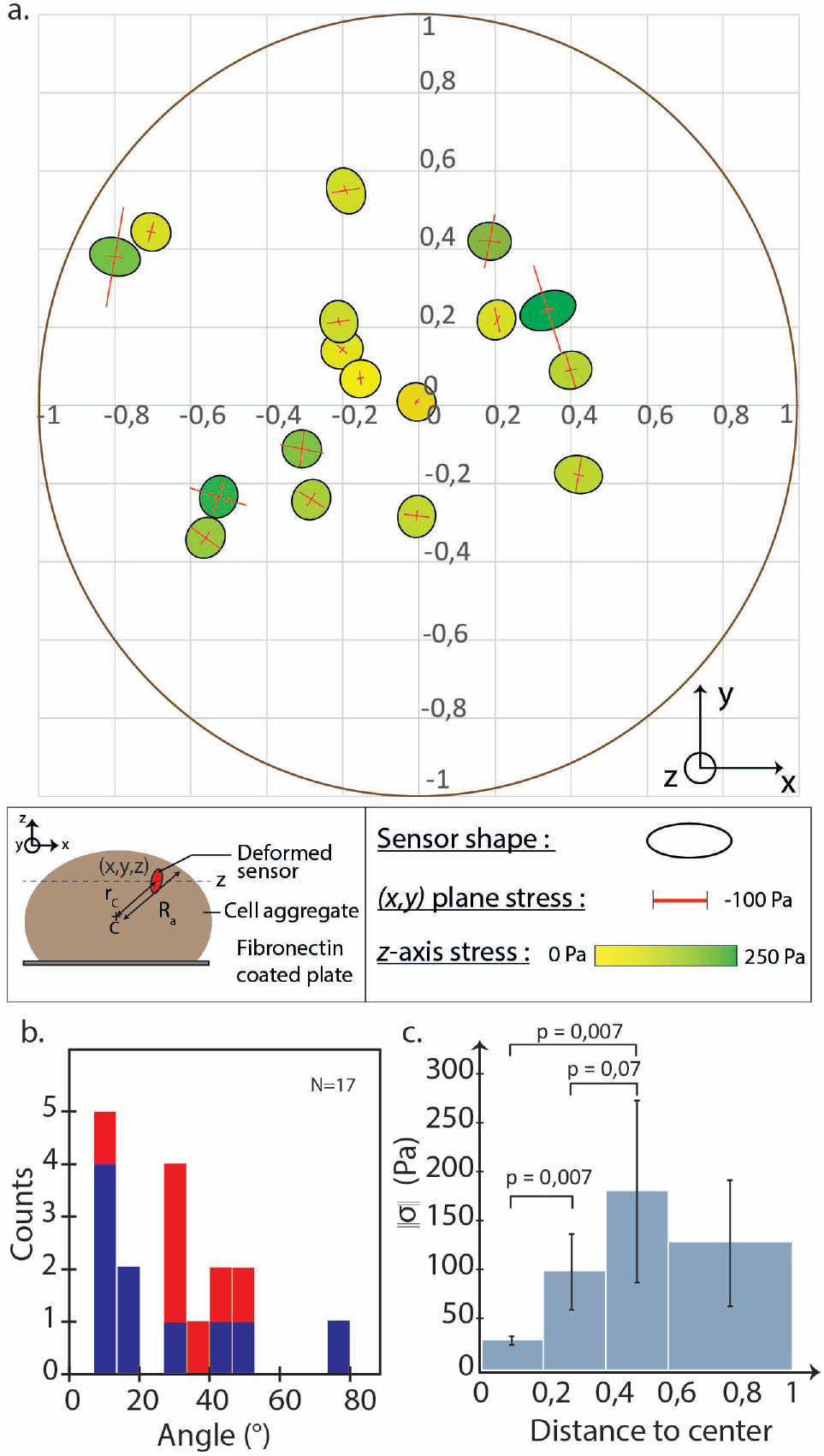
(a) Stress distribution map in CT26 aggregates, from 17 sensors inserted in 13 different aggregates. Each sensor is set at its normalized position *r* = *r*_*c*_*/R*_*a*_. Its shape projected in the (*x, y*) plane is represented by an ellipse (the ellipticity is artificially multiplied by a factor 4 to make it more easily visible). The main shear stress components in the (*x, y*) plane are shown as red bars, while the projection of the stress in the *z* direction is represented by a color code. (b) Distribution of angles between the radial direction and the sensor longer axis direction (blue: sensors showing a difference in half axes larger than the estimated accuracy 0.5 µm ; red: other sensors). The sensors are principally compressed in the orthoradial direction. (c) Histogram of the shear stress amplitude ‖*σ*‖ versus normalized distance to the center *r*.

By looking at the stress orientation and amplitude, one retrieves several pieces of information :

First, the in-plane main axes of the sensors are mostly aligned along the radial and orthoradial directions of the aggregate referential. Fig. 5b represents the distribution of angles between the radial direction and the sensor’s longer axis direction (blue: sensors showing a difference in half axes larger than the estimated accuracy threshold 0.5 µm; red: other sensors). This distribution is non-uniform and indicates that the sensors are principally compressed in the orthoradial direction.

Second, the component *σ*_*zz*_, represented by a color code in Fig.5a, is always positive and ranges between 0 and 250 Pa (see **Discussion**).

Third, the stress amplitude ‖ *σ ‖*varies from the center to the edge of the aggregate. ‖*σ ‖* is defined as the norm of the stress deviator:

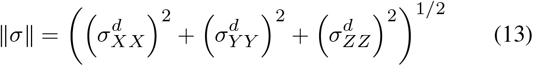

The histogram of ‖*σ*‖ is represented in Fig. 5c versus the normalized distance *r*. Despite the uncertainty, we observe a significant trend for ‖*σ*‖ to increase with *r*, and possibly to reach a maximum and decrease when approaching the edge of the aggregate.

In principle the evolution of this stress map can be followed in time. We were able to image the spreading of some aggregates deposited on the bottom plate of the Petri dish, during a few hours, by taking stacks every 15 min. In most cases, the axes orientations and half-lengths of the sensors remained stable with time, within experimental accuracy. Longer recordings would be necessary to see an evolution, and to follow the aggregate spreading process until its term.

### Stress distribution in the prechordal plate of zebrafish embryos

*In vivo*, the spatial distribution of mechanical stresses, their inhomogeneities and their local anisotropy play a determinant role in the morphogenesis process, since they directly influence cell polarization and migration. For instance, it was established *in vitro* that Xenopus prechordal plate cells can be polarized by application of a mechanical stress of a few Pa (Weber et al., 2012). The prechordal plate (PPl) is a group of cells, that are the first ones to internalize on the dorsal side of the embryo, at the onset of gastrulation. During gastrulation, they migrate in direction of the animal pole, followed by notochord precursors (Kimmel et al., 1995; Solnica-Krezel et al., 1995). Based on this observation, it was proposed that migration of the PPl is guided *in vivo* by the existence of stress anisotropies within the tissue, used by cells as directional cues (Weber et al., 2012; Behrndt and Heisenberg, 2012). Our sensors, directly measuring the 3D stress anisotropy, and allowing to map the stress in the tissue, seemed particularly well suited for such an application. Also, the prechordal plate appeared as a good model to demonstrate their *in vivo* capabilities.

#### Spatial distribution

The prechordal plate and the notochord cells are labelled in the *Tg(gsc:GFP)* line, which was used in these experiments. Sensors were implanted in the PPl of seven different embryos. Some of them could be followed over time, by taking images every 30s or 60s. An overview of the full dataset is shown in supplementary files, table S1. We report here a selection of 12 measurements, at different stages of gastrulation, from 60 to 85% of epiboly. The common effective shear modulus of all the sensors was *µ*_*d*_ = 430 Pa. To analyze the stress spatial distribution, the PPl was divided into 9 zones (front/middle/rear and left/center/right), as shown in figure 6e. For legibility, the projection of the shear stresses in the PPl (*x, y*) plane is drawn as an ellipse for each sensor, while the projection along the perpendicular axis *z* (confounded with the observation axis) is represented by a color code. In the PPl plane, *x* is the direction of the PPl progression and *y* the perpendicular one. All the stress components lie in a range comprised between +60 and − 60 Pa, with approximately equal distribution between positive and negative values.

**Fig. 6.**
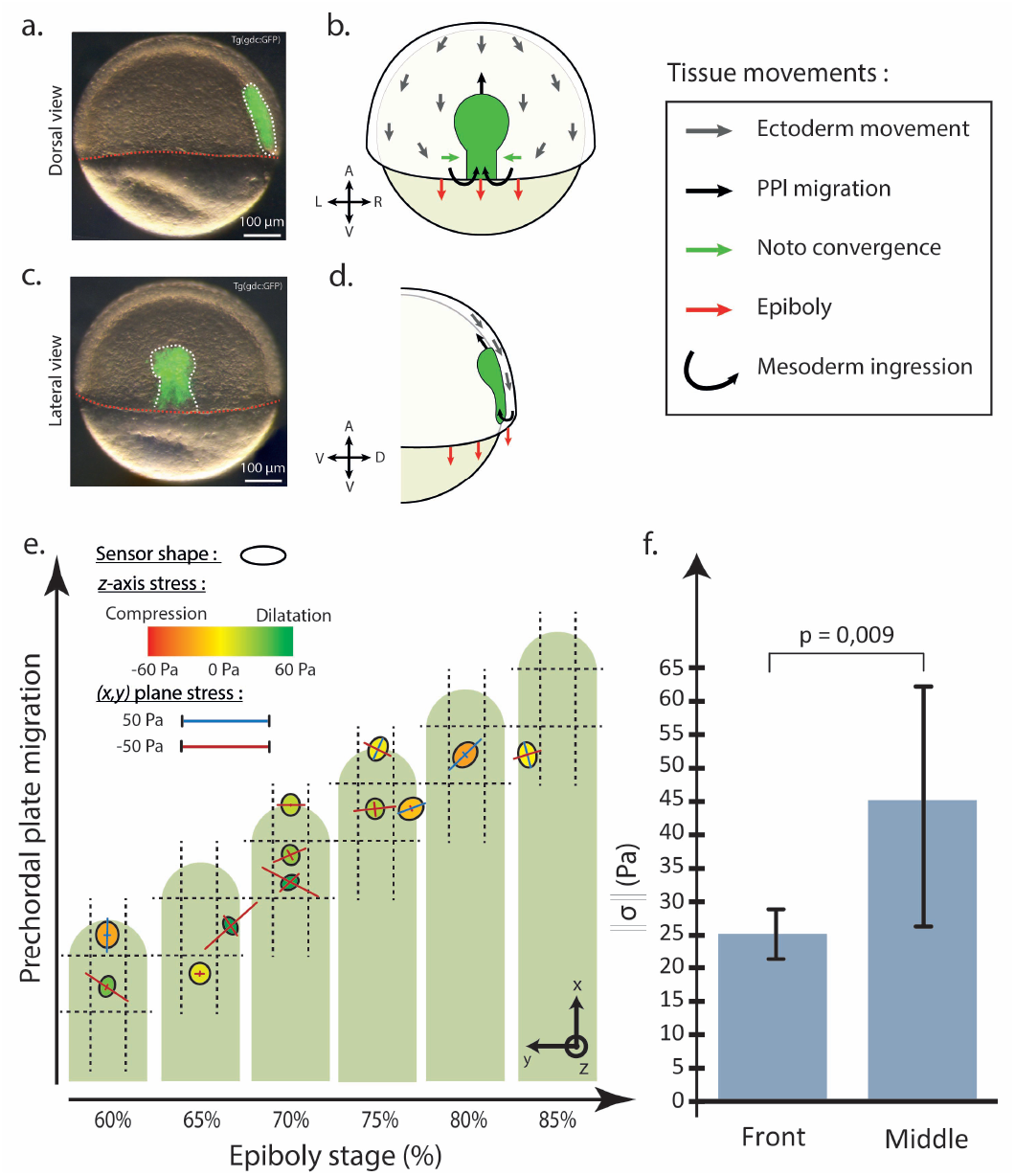
Measurements of the shear stresses in the zebrafish prechordal plate (PPl) during epiboly. (a-d) : Ectoderm, prechordal plate (PPl) and notochord precursors (Noto) movements during gastrulation. (a,c) Bright fields and fluorescence images of a zebrafish Tg(gsc:GFP) embryo at 60% epiboly. Prechordal plate and notochord cells, expressing GFP in this transgenic line, are highlighted by the white dashed line. Red dashed line marks the margin of the embryo. (b,d) Schematics of corresponding pictures showing morphogenetic movements of the different tissues during gastrulation. Crossed arrows indicate animal-vegetal (A-V), left-right (L-R) and dorsal-ventral (D-V) embryonic axes of respective view. Panels a, b, c and d have been inspired by Smutny et al. (2017). (e) Mapping of shear stresses in the PPl at different epiboly stages (12 measurements on 7 different embryos). The projection on the (*x, y*) plane of the main shear stresses is drawn as an ellipse, while the projection along the normal axis *z* is represented by a color code. (f) Comparison of the shear stress amplitude ‖*σ*‖, averaged at the front (*N* = 3) and in the middle (*N* = 7) of the PPl.

From figure 6e, no evident correlation emerges between the sensor location in one of the 9 zones of the PPl and the stress orientation and amplitude in the same zone. However, in Fig. 6f, the value of the shear stress amplitude ‖*σ*‖, averaged over the left/center/right zones and over the different epiboly stages, is compared at the front (*N* = 3) and at the middle (*N* = 7) of the plate. The difference is significant and is a first indication that stress gradients exist in the PPl.

#### Time evolution

We were able to follow the evolution of the stress components for 7 sensors, during 15 to 30 min, at different stages of epiboly.They did not show any significant changes, except for one event which we describe now.

A sensor inserted in the PPl, and migrating with it, was imaged during 15 min, at the stage 70 − 75% of epiboly. In Figure 7, the three main components *σ*_*XX*_, *σ*_*Y Y*_, and *σ*_*ZZ*_ are plotted versus time. Each principal axis (*X, Y, Z*) is color-labelled according to its nearest axis of the PPl referential: *x* (red), *y* (yellow) and *z* (blue).

**Fig. 7.**
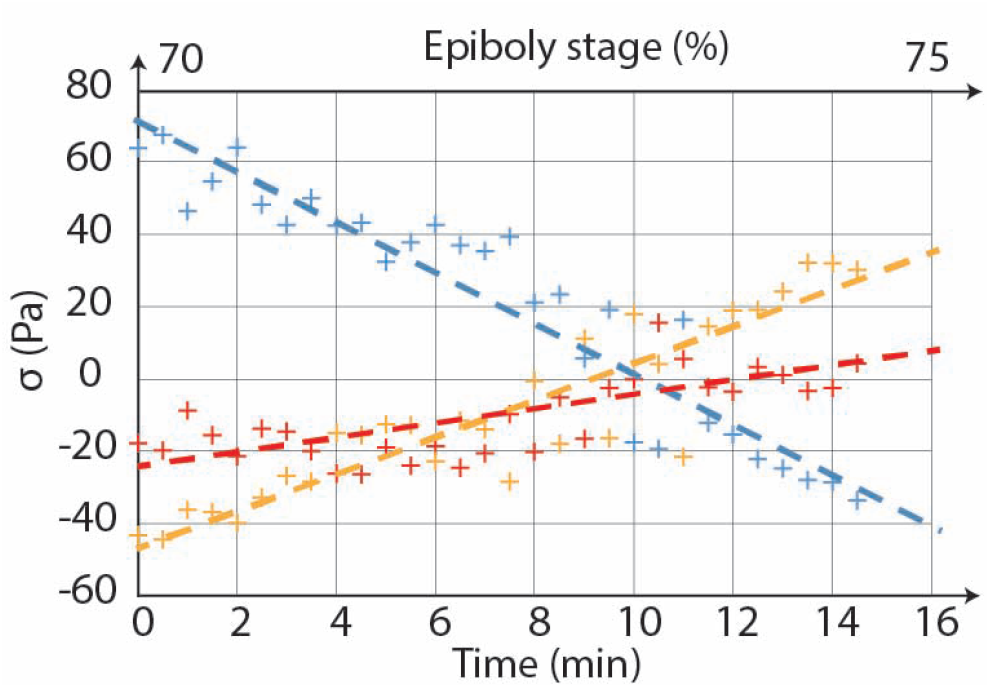
Time evolution of the main components of the shear stress tensor, for a sensor located in the central zone of the PPl of a zebrafish embryo. Each principal axis (*X, Y, Z*) is color-labelled according to its nearest axis of the PPl referential. Within 15 minutes, the stress amplitude along the principal axis closest to the *z* axis (blue) changes from positive (extension) to negative (compression). In the same time, the stress along the axis closest to the *y* axis follows the opposite evolution (yellow), and so does, to a lesser extent, the component close to *x* axis (red). Dashed lines are guides for the eye.

In Fig. 7 one can follow the evolution of each shear stress component with time. The stress amplitude along the principal axis closest to the *z* axis (blue) changes from positive (extension) to negative (compression). In the same time, the stress along the axis closest to the *y* axis follows the opposite evolution (yellow), and so does, to a lesser extent, the component close to *x* axis (red). Their directions remain stable except for some small fluctuations. This event is a clear signature of a main change in the stress partition, which takes place within a few minutes at this stage of epiboly. The short duration of this event might explain why it has been observed only once, out of 7 assays. Of course, no general conclusion can be extracted from one single event, but its occurrence demonstrates that the technique enables to follow the time evolution of the shear stress tensor during the prechordal plate migration.

## DISCUSSION

In both experiments, *in vitro* and *in vivo*, we have demonstrated that our pipeline of techniques, based on the use of PDMS elastic microsensors embedded in living tissues, can be used to locally determine the amplitudes and orientations of the shear stress components, to map them across the tissue, and to retrieve their temporal evolution.

*In vitro*, for freely spreading cell aggregates, the order of magnitude of the shear stress amplitude typically lies between 10 and 100 Pa, consistently with other measurements in similar systems (Lucio et al., 2017; Lee et al., 2019; Mohagheghian et al., 2018). Moreover, Fig. 5c shows that the deviator stress amplitude ‖*σ*‖ increases from the aggregate center to its edge, and that the stress component along the orthoradial direction is larger than along the radial one. On the other hand, according to our observations, the optical axis *Oz* systematically coincides with one of the main shear stress axis, with a positive value of the shear (extension). A possible explanation would be that, besides the applied geometrical correction due to light refraction, light diffusion in the tissue may also affect the quality of the image, especially at large depth inside the tissue (≳100 µm). If this is the case, the systematic elongation of the sensor in the *z* direction could be an artefact related to the imaging method. Fortunately, this does not affect our conclusions concerning the radial/orthoradial privileged orientations in the (*x, y*) plane, nor the variations of the stress amplitude with reduced distance *r*.

It is interesting to compare these results with those reported in (Lee et al., 2019) with hydrogel sensors embedded in spherical aggregates made of HS-5 fibroblasts. In this paper two components of the full stress tensor 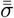 were measured *vs* the distance to the aggregate center, respectively in the radial and orthoradial directions in the observation plane. Both their amplitudes are comprised between − 400 and − 1500 Pa. Their average value represents the isotropic part of the stress : it is negative and thus corresponds to a compression. The difference between the two components, which is the local shear stress, is of the order of ± 100 Pa, comparable to their measurements accuracy. This order of magnitude is similar to ours. Moreover, they observe that both stress components increase from the aggregate edge, to reach a maximum value, and then decrease again towards the aggregate center. Although the two experiments give access to different quantities (total shear in their case, shear stress in ours), the behaviors observed in both situations are consistent.

*In vivo*, in the prechordal plate of the Zebrafish embryo, we were able to measure both the amplitude and orientation of the main stress components at different stages of epiboly. The retrieved values are of the expected order of magnitude, 10^2^ Pa, and the sensitivity is of order of 10^1^ Pa. From our data we can highlight two emerging trends, which of course will have to be confirmed by further measurements. First, following the evolution of the main stress components with time have shown that some specific events may occur, which take place within a short time interval as compared to the gastrulation, and which denote important changes in the stress partition. Such events have to be systematically identified, to determine whether they occur at some particular stages of epiboly, and to learn about their specific role in the full migration process. Secondly, the total stress amplitude appears larger at the center of the PPl than at its front, which supports the hypothesis that stress gradients exist in the PPl and might play an active role in the PPl migration. A systematic survey of the stresses amplitude and orientation in the different PPl regions is needed to comfort this assumption. Another important issue, which was not investigated here, concerns the stress distribution in the direction orthogonal to the PPl plane. Indeed, one suspects that the friction of the PPl over the neurectoderm might play an important role in the migration process (Smutny et al., 2017). Thus, further investigations will also have to include the vertical position of the sensor inside the PPl as one of the relevant parameter of the problem.

To conclude, we have assembled a pipeline of techniques which meets all the requirements to quantitatively map in 3D + time the local shear stresses in living tissues, with a sensitivity of order of 10 Pa. In addition to other complementary techniques, it appears as a valuable tool to investigate the role of mechanical constraints in morphogenesis and development.

## Acknowledgements

We acknowledge Alain Richert, Gaëtan Mary, Magid Badaoui, Alain Ponton and Sandra Lerouge for their assistance and advices in experimental handling, Hélène Delanoé-Ayari for critical reading of the manuscript, and Mathieu Roché and Julien Dervaux for fruitful discussions. We acknowledge the ImagoSeine facility, at Institut Jacques Monod, especially Xavier Baudin and Orestis Faklaris, and the Polytechnique Bioimaging Facility, for assistance with live imaging on their equipments.

## Contribution

A.S, F. Gr. and F. Ga. designed the experiments, with the help of G.C., A.A., A.B, N.D. ; A.S. carried out the experiments and data analysis, together with A.B. for the Zebrafish part under the supervision of N. D. ; G.C. supervised the chemistry of fluorescent labelling ; A.A. conducted the uniaxial rheometry ; F. Ga. did the appendix calculations ; C.N. inspired the scientific process and methodology ; F. Gr. and F. Ga. supervised the work ; A.S, F. Gr. and F. Ga. wrote the manuscript. All authors have read, corrected and approved the manuscript.

## Data availability

The python code used for active contour detection is available here : https://github.com/scikit-image/scikit-image/blob/main/skimage/segmentation/morphsnakes.py.

An example of contour determination can be found here : https://github.com/Alex-code-lab/3D_contour_detection_and_fit_by_an_ellipsoid.

## APPENDIX EXPRESSION OF THE LOCAL STRESSES IN AN AGGREGATE SUBMITTED TO AN UNIAXIAL COMPRESSION

In this section we describe a model to calculate all the components of the stress tensor at any point of an aggregate submitted to an uniaxial compression, from which we derive the expression of the deformation of an elastic incompressible sensor embedded in the aggregate.

### Notations

We assume that the aggregate, initially spherical, is squeezed between two plates applying on it a force 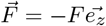, and that the problem respects the cylindrical symmetry around axis *Oz*. The notations are explicited in Fig. 8. We use cylindrical coordinates *M* (*r, ϕ, z*), *O* is the aggregate center. *R*(*z*) is the radius of the curve generating the aggregate cylindrical surface. The principal radii of curvature are:

**Fig. 8.**
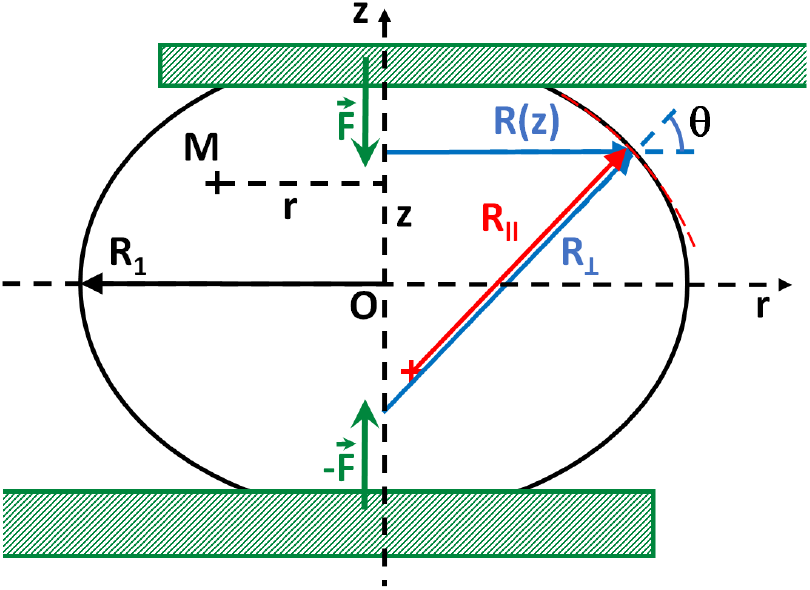
Scheme of a squeezed aggregate, assumed to respect cylindrical symmetry around *Oz*. The current point M is identified by its coordinates (*r, ϕ, z*) where *ϕ* is the azimutal angle; *R*(*z*) is the radius of the curve generating the aggregate cylindrical surface; *R*_⊥_ and *R*_‖_ are the principal curvature radii; *R*_1_ is the equatorial value of 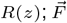 is the force exerted by the plates on the aggregate.

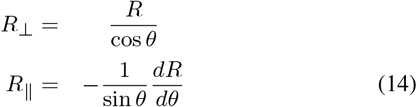

with 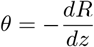.

In the particular case of a circular profile, which is a non-necessary but sufficient approximation for most situations, one has:

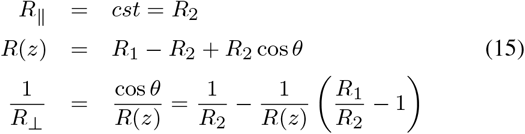

where *R*_1_ is the equatorial radius.

The local stress tensor is written as :

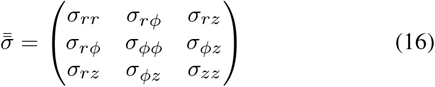

### Mechanical equilibrium

In the absence of external volumic force, the mechanical equilibrium condition is written : 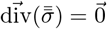, *i*.*e*. in cylindrical coordinates:

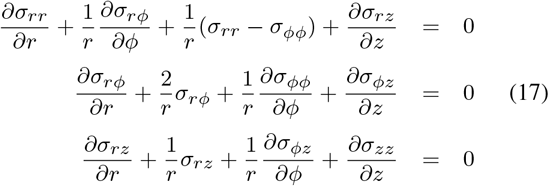

Due to the cylindrical symmetry, all the stress components are independent on the azimutal angle *ϕ*. Furthermore *σ*_*rϕ*_ = *σ*_*zϕ*_ = 0. Thus the above equations simplify into:

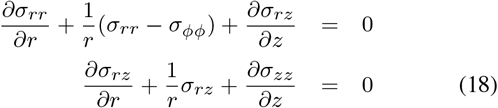

We can also write the mechanical equilibrium condition at the aggregate surface, *i*.*e*. for *r* = *R*(*z*). By equilibrating the local stresses at the boundary with the external pressure *p*_*a*_ outside the aggregate and the Laplace pressure, one finds in projection along the *r* and *z* axes at any point *M* (*R, z*) of the surface:

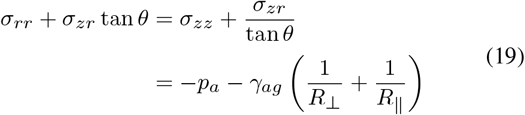

We recall that *γ*_*ag*_ represents the surface tension between the aggregate and the external medium.

Finally, following Norotte et al. (2008), the global balance of forces exerted on the aggregate, in a plane perpendicular to *Oz*, at coordinate *z*, can be expressed as:

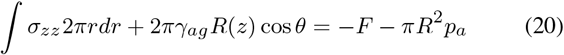

### Expression of 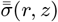

We assume in the following that the expressions of the components *σ*_*zz*_ (*r, z*), *σ*_*rr*_(*r, z*) and *σ*_*ϕϕ*_(*r, z*) can be approximated by a Taylor expansion to order *r*^2^, according to :

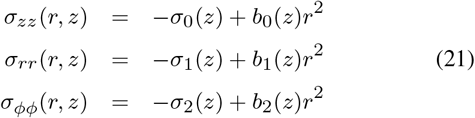

The functions *σ*_*i*_(*z*) and *b*_*i*_(*z*) (*i* = 0, 1, 2) account for the *z*-dependence of the stress components. As shown below, they can be explicitly calculated from Eqs. (18) to (20). Note that *σ*_*i*_(*z*) is positive in case of a compression.

From equation (18) one derives the expression of *σ*_*rz*_ :

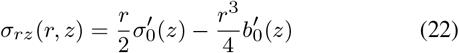

where the prime stands for the *z*-derivative.

Introducing the expression of *σ*_*rz*_ (Eq.(22)) in Eq. (18) yields:

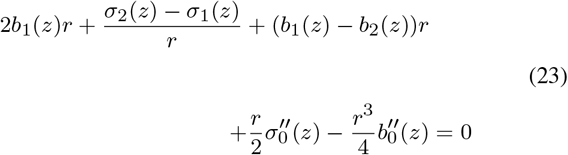

Since the components of 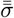 must not diverge in *r* = 0, necessarily *σ*_2_(*z*) = *σ*_1_(*z*). There remains :

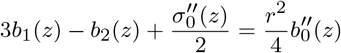

Because we limit the Taylor development to order *r*^2^ for *σ*_*rr*_ and *σ*_*ϕϕ*_, *b*_1_ and *b*_2_ do not depend on *r*. Consequently 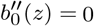 and thus *b*_0_(*z*) = *Az* + *B*, where *A* and *B* are two constants. Moreover, by *z* to − *z* symmetry, *σ*_*zz*_ must be an even function of *z*, which implies *A* = 0. Therefore:

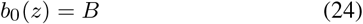

Here *B* is a constant independent of *z*, but it may depend on time *t* if the stresses in the aggregate evolve with time.

At this stage an additional relation between *b*_1_(*z*) and *b*_2_(*z*) is required to close the equation system (18) to (20) and complete the calculation. In the following, we will assume for simplicity that the stress projection in any plane orthogonal to *Oz* is isotropic, which means *σ*_*rr*_(*r, z*) = *σ*_*ϕϕ*_(*r, z*) and thus *b*_1_(*z*) = *b*_2_(*z*). This most simple hypothesis can be partially justified by geometrical arguments, related to the incompressibility of the material, which we will not develop here. Other assumptions remain possible, but we have checked, after performing the whole calculation in different cases, that the result is not modified except for some minor numerical factors. Under this assumption, Eq. (23) simplifies into:

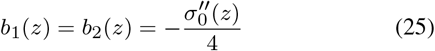

The integral of Eq. (20) may be calculated using Eq. (21), which leads to :

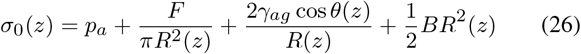

Eqs. (24), (25) and (26) are sufficient to calculate all the components of the stress tensor, at any point of a cylindrical aggregate of generator *R*(*z*). For instance, one finds for *σ*_*zz*_ (*r, z*) :

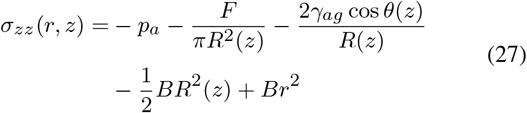

The only remaining free parameter in Eq. (27) is the constant *B*, the value of which will be discussed at the end of this section.

The other components *σ*_*zr*_(*r, z*) and *σ*_*rr*_(*r, z*) = *σ*_*ϕϕ*_(*r, z*) can also be written after some long but straightforward calculations, using Eqs. (21), (22) and the boundary condition Eq. (19) to calculate *σ*_1_(*z*).

In the following, we only focus on the case of an aggregate with a circular profile, *i*.*e. R*_*‖*_ = *R*_2_ = *cst*, and we write down the stress components in its equatorial plane *z* = 0. In this plane, *R*_⊥_ = *R*_1_ and *θ* = 0, which leads to :

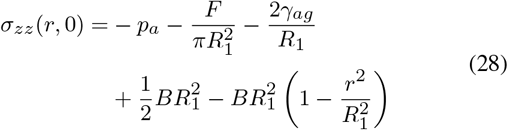

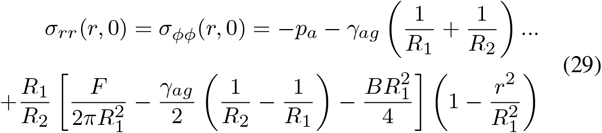

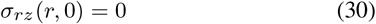

### Deformation of a sensor embedded in the aggregate

We consider an elastic incompressible sensor, spherical at rest, embedded in the aggregate, of effective shear modulus *µ*_*e*_. We assume that the mechanical properties of the sensor and of the aggregate are similar, and that the sensor diameter is small compared to the aggregate’s one, so that the sensor inclusion does not perturbate the stress distribution in the aggregate. Due to incompressibility, only shear deformations of the sensor are admitted, and the the strain-stress tensors relationship reduces to (see Eq. (2)):

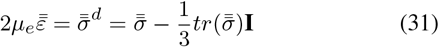

In the following, we assume that the sensor size is negligible with respect to the aggregate size. We also assume for simplicity that it is located in the equatorial plane *z* = 0, although the calculations could in principle be performed for any position in the aggregate. We introduce 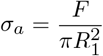 as the average stress exerted by the force *F* on the equatorial plane. Using Eqs. (28) and (29), one finds:

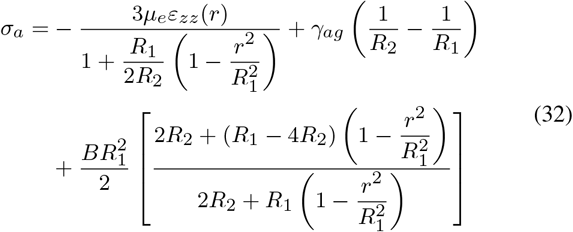

Equivalently, in Eq. (32) one can replace −*ε*_*zz*_ by +2*ε*_*rr*_, since incompressibility implies 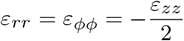.

At the aggregate center (*r* = 0, *z* = 0), Eq. (32) simplifies into:

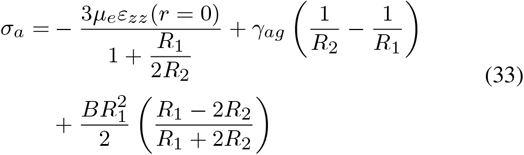

At the aggregate edge (*r* = *R*_1_, *z* = 0), one has :

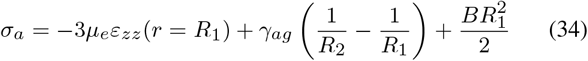

### Choice of the value of *B*

In this model, *B* has a homogeneous value over the volume of the sensor, but it may vary over time. Indeed, in the experiment one applies at *t* = 0 a step of displacement to the rigid plate, which imprints a constant deformation to the aggregate. Immediately after the step, the stress in the aggregate is inhomogeneous, but it rapidly evolves, within a few minutes, through different relaxation mechanisms, to become homogeneous again at the end of relaxation. *B*(*t*) is therefore a function of time which must relax towards zero at infinite time. To interpret our data concerning *σ*_*a*_(*ε*), we discard the first moments of the relaxation, and we only consider the long time limit, for which we expect the stresses to be homogeneous again, allowing us to make the approximation *B* = 0. Within this assumption, Eq. (33) exactly reduces to Eq. (12) used in the main text. To comfort this assumption, we also performed the data analysis by taking a non-zero value for *B*, leading to parabolic variations for *σ*_*zz*_ as a function of *r*. Assuming for instance that *σ*_*zz*_ vanishes at the aggregate edge (*r* = *R*_1_, *z* = 0), from Fig. (4c) one retrieves *µ*_*e*_ = 670 Pa, instead of *µ*_*e*_ = 790 Pa in the case *B* = 0. Considering all other sources of uncertainty, the results do not appear significantly different.

## REFERENCES

S. Angeli and T. Stylianopoulos. Biphasic modeling of brain tumor biomechanics and response to radiation treatment. Journal of Biomechanics, 49 (6):1524–1531, 2016.

K. Bambardekar, R. Clément, O. Blanc, C. Chardès, and P.-F. Lenne. Direct laser manipulation reveals the mechanics of cell contacts in vivo. Proceedings of the National Academy of Sciences, 112(5):1416–1421, 2015.

M. Behrndt and C.-P. Heisenberg. Spurred by resistance: Mechanosensation in collective migration. Developmental Cell, 22(1):3–4, 2012.

M. Bendaoud. Développement de méthodes d’extraction de contours sur des images à niveaux de gris. PhD thesis, 2017.

J. Bico, Reyssat, and B. Roman. Elastocapillarity: When surface tension deforms elastic solids. Annual Review of Fluid Mechanics, 50(1):629–659, 2018.

I. Bonnet, P. Marcq, F. Bosveld, L. Fetler, Y. Bellaiche, and F. Graner. Mechanical state, material properties and continuous description of an epithelial tissue. Journal of The Royal Society Interface, 9(75):2614–2623, 2012.

N. Borghi, M. Sorokina, O. Shcherbakova, W. Weis, B. Pruitt, W. Nelson, and A. Dunn. E-cadherin is under constitutive actomyosin-generated tension that is increased at cell-cell contacts upon externally applied stretch. Proceedings of the National Academy of Sciences, 109(31):12568–12573, 2012.

A. Boutillon, F. A. Giger, and N. B. David. Analysis of In Vivo Cell Migration in Mosaic Zebrafish Embryos, pages 213–226. Springer New York, New York, NY, 2018.

G. Brodland, J. Veldhuis, S. Kim, M. Perrone, D. Mashburn, and M. S. Hutson. Cellfit: A cellular force-inference toolkit using curvilinear cell boundaries. PLoS ONE, 9(6):1–15, 2014.

H.-J. Butt, B. Cappella, and M. Kappl. Force measurements with the atomic force microscope: Technique, interpretation and applications. Surface Science Reports, 59(1-6):1–152, 2005.

O. Campàs. A toolbox to explore the mechanics of living embryonic tissues. Seminars in Cell and Developmental Biology, 55:119–130, 2016.

O. Campàs, T. Mammoto, S. Hasso, R. Sperling, D. O’Connell, A. Bischof, and D. Ingber. Quantifying cell-generated mechanical forces within living embryonic tissues. Nature Methods, 11(2):183–189, 2014.

A. Carbonaro, K.-N. Chagua-Encarnacion, C.-A. Charles, T. Phou, C. Ligoure, S. Mora, and D. Truzzolillo. Spinning elastic beads: a route for simultaneous measurements of the shear modulus and the interfacial energy of soft materials. Soft Matter, 16:8412–8421, 2020.

V. Caselles, F. Catte, B. Coll, and F. Dibos. A geometric model for edge detection. Numerische Mathematik, 66(1):1–31, 1993.

K. Chiou, L. Hufnagel, and B. Shraiman. Mechanical stress inference for two dimensional cell arrays. PLoS Computational Biology, 8(5):1–9, 2012.

M. Delarue, F. Montel, D. Vignjevic, J. Prost, J. Joanny, and G. Cappello. Compressive stress inhibits proliferation in tumor spheroids through a volume limitation. Biophysical Journal, 107:1821–1828, 2014.

N. Desprat, A. Richert, J. Simeon, and A. Asnacios. Creep function of a single living cell. Biophysical Journal, 88(3):2224–2233, 2005.

N. Desprat, A. Guiroy, and A. Asnacios. Microplates-based rheometer for a single living cell. Reviex of scientifc intruments, 77:055111, 2006.

N. Desprat, W. Supatto, P.-A. Pouille, E. Beaurepaire, and E. Farge. Tissue deformation modulates twist expression to determine anterior midgut differentiation in drosophila embryos. Developmental Cell, 15(3):470–477, 2008.

M. Doitsidou, M. Reichman-Fried, J. Stebler, M. Köprunner, J. Dorries, D. Meyer, C. Esguerra, T. Leung, and E. Raz. Guidance of primordial germ cell migration by the chemokine sdf-1. Cell, 11(5):647–659, 2002.

M. Dolega, M. Delarue, F. Ingremeau, J. Prost, A. Delon, and G. Cappello. Cell-like pressure sensors reveal increase of mechanical stress towards the core of multicellular spheroids under compression. Nature Communications, 8(4):14056, 2017.

B. Elkin, E. Azeloglu, K. Costa, and B. Morrison III. Mechanical heterogeneity of the rat hippocampus measured by atomic force microscope indentation. Journal of Neurotrauma, 24(5):812–822, 2007.

K. Franze. Atomic force microscopy and its contribution to understanding the development of the nervous system. Current Opinion in Genetics and Development, 21(5):530–537, 2011.

M. Gomez-Gonzalez, E. Latorre, M. Arroyo, and X. Trepat. Measuring mechanical stress in living tissues. Nature Reviews Physics, 2(6):300–317, 2020.

C. Grashoff, B. Hoffman, M. Brenner, R. Zhou, M. Parsons, M. Yang, and M. Schwartz. Measuring mechanical tension across vinculin reveals regulation of focal adhesion dynamics. Nature, 466(7303):263–266, 2010.

K. Guevorkian, M.-J. Colbert, M. Durth, S. Dufour, and F. Brochard-Wyart. Aspiration of biological viscoelastic drops. Physical Review Letters, 104 (21):218101, 2010.

B. Guirao, S. U. Rigaud, F. Bosveld, A. Bailles, J. Lopez-Gay, S. Ishihara, K. Sugimura, F. Graner, and Y. Bellaiche. Unified quantitative characterization of epithelial tissue development. eLife, 4:e08519, 2015.

K. Haejune. Controlled production of emulsion drops using an electric field in a flow-focusing microfluidic device. Applied Physics Letters, 91(13):211–218, 2007.

H. Hamada. Role of physical forces in embryonic development. Seminars in Cell and Developmental Biology, 47-48:88–91, 2015.

C. Heisenberg and Y. Bellaïche. Forces in tissue morphogenesis and patterning. Cell, 153:948–962, 2013.

W. Helfrich. Elastic properties of lipid bilayers : theory and possible experiments. Z Naturforsch, 28(11):693–703, 1973.

R. Herrera-Perez and K. Karen. Biophysical control of the cell rearrangements and cell shape changes that build epithelial tissues. Current Opinion in Genetics and Development, 51:88–95, 2018.

R. Hiramatsu, T. Matsuoka, C. Kimura-Yoshida, S.-W. Han, K. Mochida, T. Adachi, and I. Matsuo. External mechanical cues triggerthe establishment of the anterior-posterioraxis in early mouse embryos. Developmental Cell, 27(2):131–144, 2013.

E. K. Hobbie, S. Lin-Gibson, and S. Kumar. Non-brownian microrheology of a fluid-gel interface. Physical Review Letters, 100(7):076001, 2008.

R. Hochmuth. Micropipette aspiration of living cells. Journal of Biomechanics, 44(1):15–22, 2000.

B. G. Hosu, K. Jakab, P. Bánki, F. I. Tóth, and G. Forgacs. Magnetic tweezers for intracellular applications. Review of Scientific Instruments, 74(9): 4158–4163, 2003.

S. Ishihara, K. Sugimura, S. Cox, I. Bonnet, Y. Bellaïche, and F. Graner. Comparative study of non-invasive force and stress inference methods in tissue. The European Physical Journal E, 36(4):45, 2013.

M. Kass, A. Witkin, and D. Terzopoulos. Snakes : Active contour models. International Journal of Computer Vision, 1:321–331, 1988.

M. Krieg, Y. Arboleda-Estudillo, P.-H. Puech, J. Käfer, F. Graner, D. Müller, and C.-P. Heisenberg. Tensile forces govern germ-layer organization in zebrafish. Nature Cell biology, 10(4):429–436, 2008.

L. Landau, E. Lifshitz, A. Kosevich, and L. Pitaevskii. Theory of Elasticity : course of theoretical physics, volume 7. Elsevier, 1986.

K. Lau, H. Tao, H. Liu, J. Wen, K. Sturgeon, N. Sorfazlian, and S. Hopyan. Anisotropic stress orients remodelling of mammalian limb bud ectoderm. Nature Cell Biology, 17(5):569–579, 2015.

L. Le Goff, H. Rouault, and T. Lecuit. A global pattern of mechanical stress polarizes cell divisions and cell shape in the growing drosophila wing disc. Development, 140(19):4051–4059, 2013.

W. Lee, N. Kalashnikov, S. Mok, R. Halaoui, E. Kuzmin, A. Putnam, and C. Moraes. Dispersible hydrogel force sensors reveal patterns of solid mechanical stress in multicellular spheroid cultures. Soft Matter, 13(23): 4210–4213, 2019.

A. A. Lucio, A. Mongera, E. Shelton, R. Chen, A. M. Doyle, and O. Campas. Spatiotemporal variation of endogenous cell-generated stresses within 3D multicellular spheroids. Scientific Reports, 7:12022, 2017.

P. Marquez-Neila, L. Baumela, and L. Alvarez. A morphological approach to curvature-based evolution of curves and surfaces. IEEE Transactions on Pattern Analysis and Machine Intelligence, 36(1):2–17, 2014.

F. Mazuel, M. Reffay, V. Du, J.-C. Bacri, J.-P. Rieu, and C. Wilhelm. Magnetic flattening of stem-cell spheroids indicates a size-dependent elastocapillary transition. Physical Review Letters, 114(9):1416–1421, 2015.

A. Mgharbel, H. Delanoë-Ayari, and J. Rieu. Measuring accurately liquid and tissue surface tension with a compression plate tensiometer. HFSP Journal, 3(3):213–221, 2009.

J. Mitchison and M. Swann. The mechanical properties of the cell surface : I. the cell elastimeter. Journal of Experimental Biology, 31(3):443–460, 1954.

D. Mitrossilis, J. Fouchard, A. Guiroy, N. Desprat, N. Rodriguez, B. Fabry, and A. Asnacios. Single-cell response to stiffness exhibits muscle-like behavior. Proceedings of the National Academy of Sciences, 106(43):18243–18248, 2009.

D. Mitrossilis, J. Fouchard, D. Pereira, F. Postic, A. Richert, M. Saint-Jean, and A. Asnacios. Real-time single-cell response to stiffness. Proceedings of the National Academy of Sciences, 107(38):16518–16523, 2010.

E. Mohagheghian, J. Luo, J. Chen, G. Chaudhary, J. Chen, J. Sun, and N. Wang. Quantifying compressive forces between living cell layers and within tissues using elastic round microgels. Nature Communications, 9(1):1878, 2018.

A. Mongera, P. Rowghanian, H. Gustafson, E. Shelton, D. Kealhofer, E. K. Carn, and O. Campàs. A fluid-to-solid jamming transition underlies vertebrate body axis elongation. Nature, 561(7723):401–405, 2018.

S. Monnier, M. Delarue, B. Brunel, M. Dolega, A. Delon, and G. Cappello. Effect of an osmotic stress on multicellular aggregates. Methods, 94:114– 119, 2016.

K. C. Neuman and A. Nagy. Single-molecule force spectroscopy: optical tweezers, magnetic tweezers and atomic force microscopy. Nature Methods, 5(6):491–505, 2008.

U. Nienhaus, T. Aegerter-Wilmsen, and C. M. Aegerter. Determination of mechanical stress distribution in drosophila wing discs using photoelasticity. Mechanisms of Development, 126(11):942–949, 2009.

V. Nier, S. Jain, C. T. Lim, S. Ishihara, B. Ladoux, and P. Marcq. Inference of internal stress in a cell monolayer. Biophysical Journal, 110(7):1625–1635, 2016.

C. Norotte, F. Marga, A. Neagu, I. Kosztin, and G. Forgacs. Experimental evaluation of apparent tissue surface tension based on the exact solution of the laplace equation. Europhysics Letters, 81(4):46003, 2008.

J. M. Northcott, I. S. Dean, J. K. Mouw, and V. M. Weaver. Feeling stress: The mechanics of cancer progression and aggression. Frontiers in Cell and Developmental Biology, 6:17, 2018.

S. Porazinski, H. Wang, Y. Asaoka, M. Behrndt, T. Miyamoto, H. Morita, and M. Furutani-Seiki. Yap is essential for tissue tension to ensure vertebrate 3d body shape. Nature, 521(7551):217–221, 2015.

M. Rauzi, P. Verant, T. Lecuit, and P.-F. Lenne. Nature and anisotropy of cortical forces orienting drosophila tissue morphogenesis. Nature Cell Biology, 10(12):1401–1410, 2008.

P. Roca-Cusachs, V. Conte, and X. Trepat. Quantifying forces in cell biology. Nature Cell Biology, 19(7):742–751, 2017.

C. Roffay, C.-J. Chan, B. Guirao, T. Hiiragi, and F. Graner. Inferring cell junction tension and pressure from cell geometry. Develoment, 148:1–16, 2021.

T. Schluck and C. M. Aegerter. Photo-elastic properties of the wing imaginal disc of drosophila. Eur. Phys. J. E, 33(2):111–115, 2010.

M. Smutny, Z. Akos, S. Grigolon, S. Shamipour, V. Ruprecht, D. Capek, M. Behrndt, E. Papusheva, M. Tada, B. Hof, T. Vicsek, G. Salbreux, and C.-P. Heisenberg. Friction forces position the neural anlage. Nature Cell Biology, 19(4):306, 2017.

A. Souchaud. Cartographie des contraintes mécaniques in situ dans les tissus vivants. PhD thesis, Université de Paris, 2020.

R. W. Style, A. Jagota, C.-Y. Hui, and E. R. Dufresne. Elastocapillarity: Surface tension and the mechanics of soft solids. Annual Review of Condensed Matter Physics, 8(1):99–118, 2017.

K. Sugimura, P.-F. Lenne, and F. Graner. Measuring forces and stresses in situ in living tissues. Development, 143(2):186–196, 2016.

M. Tanase, N. Biais, and M. Sheetz. Magnetic tweezers in cell biology. Cell Mechanics, 22(1):473–493, 2007.

J.-Y. Tinevez, U. Schulze, G. Salbreux, J. Roensch, J.-F. Joanny, and E. Paluch. Role of cortical tension in bleb growth. Proceedings of the National Academy of Sciences, 106(44):18581–18586, 2009.

N. Träber, K. Uhlmann, S. Girardo, G. Kesavan, K. Wagner, J. Friedrichs, and J. Guck. Polyacrylamide bead sensors for in vivo quantification of cell-scale stress in zebrafish development. Scientific Reports, 9(1):17031, 2019.

M. Von Dassow, J. A. Strother, and L. A. Davidson. Surprisingly simple mechanical behavior of a complex embryonic tissue. PLoS ONE, 5(12): 958–966, 2010.

G. Weber, M. Bjerke, and D. DeSimone. A mechanoresponsive cadherin-keratin complex directs polarized protrusive behavior and collective cell migration. Developmental Cell, 22(1):104–115, 2012.

R. Wells. Tissue mechanics and fibrosis. Biochimica et Biophysica Acta : Molecular Basis of Disease, 1832(7):884–890, 2013.

Y. Xiong, A. Lee, D. Suter, and G. Lee. Topography and nanomechanics of live neuronal growth cones analyzed by atomic force microscopy. Biophysical Journal, 96(12):5060–5072, 2009.

